# TERT interacts with TFAM to activate mitochondrial fragmentation and auto/mitophagy

**DOI:** 10.64898/2026.09.18.752609

**Authors:** Erica Rossi, Jessica Marinaccio, Ilaria Festo, Ion Udroiu, Emanuela Micheli, Giulia Bertolin, Antonella Sgura

**Affiliations:** Department of Science, University “Roma Tre”, 00146, Rome, Italy; CNRS, Univ Rennes, IGDR (Institut de Genetique et Developpement de Rennes) - UMR 6290, F-35000 Rennes, France

**Keywords:** TERT, mitochondria, mitochondrial transcription factor A, interactors, mitophagy flux, mitochondrial complex IV

## Abstract

Previous studies have demonstrated that Telomerase Reverse Transcriptase (TERT) is localized inside mitochondria, where it associates with mitochondrial DNA (mtDNA), thereby influencing replication and transcription. Using a combination of quantitative fluorescence microscopy and biochemistry, we demonstrate that TERT binds indirectly to mtDNA through the interaction with mitochondrial transcription factor A (TFAM) to induce mitochondrial fragmentation. These findings reveal a novel mitochondrial interaction involving TERT and highlight a role for TERT and TFAM in the regulation of auto/mitophagy. This interaction is an important factor for mitochondrial function and contributes to the remodelling of mitochondrial respiratory capacity.

**Highlights:**

- TERT predominantly binds TFAM inside mitochondria.
- TERT induces mitochondrial fission.
- TERT/TFAM cooperates to enhance auto/mitophagy.
- TERT-overexpression promotes remodelling of the electron transport chain.

## 1. Introduction

TERT (Telomerase reverse transcriptase) is the catalytic subunit of telomerase enzyme with reverse transcriptase activity, which works together with TERC (telomerase RNA component) to elongate telomeres (Greider and Blackburn, 1989). Beyond its canonical role, TERT is able to localize inside mitochondria, thanks to the mitochondrial targeting sequence (MTS), composed by 20 amino acid residues, at its N-terminus (Santos et al., 2004a). TERT is imported through the outer and inner mitochondrial membrane translocases and reaches the mitochondrial matrix (Haendeler et al., 2009). Although several studies have shown the presence of TERT inside mitochondria under oxidative stress conditions (Haendeler et al., 2009; Marinaccio et al., 2023; Santos et al., 2004), it has been recently demonstrated that TERT is imported into mitochondria under basal conditions as well (Marinaccio et al., 2025).

Mitochondria are organelles with their own genome of 16.6 Kb, with the heavy (H) and light (L) strands both encoding genes, and a non-coding region called the D-loop. Mitochondrial DNA (mtDNA) is packaged into nucleoprotein structures called nucleoids, and one of the most extensively characterized proteins involved in this packaging is the mitochondrial transcription factor A (TFAM). TFAM is known to cooperate with a machinery of proteins required for mtDNA maintenance, such as mitochondrial single-stranded DNA binding proteins (mtSSB), DNA polymerase ϒ (POLG), and the helicase TWINKLE (Falkenberg, 2018). Furthermore, TFAM is involved in mtDNA transcription. TFAM forms an assembly to initiate transcription through its interaction with mitochondrial RNA polymerase (POLRMT) and the mitochondrial transcription factor B2 (TFB2M) (Falkenberg et al., 2024). The mitochondrial transcription initiation complex was shown to contribute to primer synthesis for mtDNA replication, therefore enabling the POLG to proceed with strand elongation. Accordingly, mtDNA replication and transcription are two closely interconnected processes (Falkenberg, 2018; Falkenberg et al., 2024).

Mitochondria have key roles in energy production and regulate cellular homeostasis and survival (Spinelli and Haigis, 2018). Mitochondrial homeostasis is maintained through a balance between mitochondrial biogenesis and clearance of damaged mitochondria (Ploumi et al., 2017). Mitochondrial biogenesis is a tightly regulated mechanism by which cells increase mitochondrial mass generating new functional mitochondria. This is based on the coordinated regulation of several processes, including the synthesis of mtDNA-encoded proteins, the import and synthesis of nuclear-encoded ones, and the replication of mtDNA (Wenz, 2013). On the other hand, cells employ an efficient clearance mechanism to remove damaged mitochondria through mitophagy. This is a specific form of autophagy that targets mitochondria for degradation through a receptor-mediated mechanism. Mitophagy receptors typically contain LC3-Interacting Region (LIR), which mediate the interaction of the key autophagy mediator LC3 with the phagophore (Ploumi et al., 2017).

Previously, Marinaccio et al. (2025) observed that TERT is associated with mtDNA without a specific binding sequence, and this to increase mtDNA replication and transcription. Additionally, it has been shown that TERT decreases mitochondrial mass without affecting mtDNA copy number. Moreover, an increased amount of smaller-sized mitochondria and of mito/autophagosomes were observed upon TERT expression (Marinaccio et al., 2025). Based on these findings, it remains to be established whether TERT binds mtDNA directly or if the binding is mediated by other nucleoid complex proteins responsible for mtDNA replication and transcription. In addition, it is still an open question whether fragmented mitochondria are organelles actively undergoing mitophagy, and whether TERT is directly involved in this process through the interaction with specific mitochondrial partners.

Here, we describe the interaction of TERT with nucleoid complex proteins, and that TERT expression drives Drp1-mediated mitochondrial fission. Furthermore, we show that the TERT/TFAM interaction is required for mitophagy activation upon TERT overexpression in U2OS cells, and for lowering the abundance of the mitochondrially-encoded Complex IV subunit COX-I. We thus propose that the TERT/TFAM interaction is a key factor for mitochondrial function, and a step in the lowering of mitochondrial respiratory capacity previously reported upon TERT expression (Marinaccio et al., 2025).

## 2. Methods

### 2.1. Cell culture and transfection

Human Colorectal Carcinoma HCT-116 (ECACC, Salisbury, UK), Osteosarcoma cell line U2OS (ECACC) and U2OS-TERT-HA obtained by retroviral transduction as previously reported (Marinaccio et al., 2025), were grown in Dulbecco’s Modified Eagle Medium High Glucose (Euroclone, Milan, Italy). The media was supplemented with 10% fetal bovine serum (Euroclone), 1000 units/ml penicillin, 10 mg/ml streptomycin and 2 mM L-glutamine (Euroclone). Cells were maintained in a humidified incubator at 37°C with 95% relative humidity and 5% CO_2_.

To study the auto/mitophagy flux, U2OS and U2OS-TERT-HA cell lines were seeded at 70% confluence in 4 well Labteks (Nunc). 24 hours after seeding, the cells were transfected with the following plasmids: pCMV Aquamarine-LC3B (Addgene plasmid n. 228559), pCMV Aquamarine-LC3B-tdLanYFP (Addgene plasmid n. 228560) and pCMV Aquamarine-LC3B-G120A-tdLanYFP (Addgene plasmid n. 228561) and co-transfected with siRNA. Negative control siRNA, SCR (AM4641) from Invitrogen (Carlsbad, CA, USA), Allstar Negative control (1027281) from Qiagen (Redwood City, CA, USA) and the *TFAM* siRNA (SI04988494) from Qiagen (USA) were used. Plasmid DNA transfection or plasmid DNA and siRNA co-transfection experiments were performed using Lipofectamine 2000 (Invitrogen, USA). For transfection with siRNA alone, Lipofectamine RNAiMAX (Invitrogen, USA) was used.

### 2.2. *in situ* Proximity Ligation Assay (*is*PLA)

To quantify the mitochondrial protein interactors of TERT with the nucleoid complex proteins, an *in situ* Proximity Ligation Assay (*is*PLA) was performed. 1×10^5^ U2OS-TERT-HA and HCT-116 cells were seeded in a 35mm petri dish, fixed with 4% paraformaldehyde and permeabilized with 0.2% PBS/Triton. In both cell lines, *is*PLA was performed using the NaveniFlex Cell MR Atto 647N PLA kit (Navinci Uppsala, Sweden) according to manufacturer’s instructions. Primary antibodies were used in pairs to detect the interactions, as follows: (I) rabbit anti-hTERT (Rockland, Limerick, PA, USA)/ mouse anti-mtRPOL (Santa Cruz Biotechnology, Dallas, TX, USA); (II) rabbit anti-hTERT/mouse anti-TWINKLE (Santa Cruz Biotechnology); (III) rabbit anti-hTERT/mouse anti-TFB2M (Santa Cruz Biotechnology); (IV) rabbit anti-hTERT/mouse anti-TFAM (Santa Cruz Biotechnology). A single primary antibody (rabbit anti-hTERT) was used as negative control of the specificity of the interaction. DNA was counterstained by DAPI (4’-6-diamidin-2-phenylindole) (Sigma-Aldrich, Merck, Darmstadt, Germany) and Vectashield anti-fade (Vector Laboratories, Burlingame, CA, USA). Images were acquired by Nikon A1 microscope (Nikon instruments Inc., Tokyo; Japan) at a 40x magnification equipped with CLSM. The analysis was performed on a total of 500 cells per condition and the summary result of PLA puncta/cells was reported.

### 2.3. Immunofluorescence, FRET/FLIM analysis and confocal microscopy

To analyze TERT-interacting proteins, 5×10^4^ of U2OS-TERT-HA cells were seeded on 15 mm round coverslips placed onto 24-well plates, fixed in 4% paraformaldehyde and permeabilized in 0.2% PBS/Triton. Cells were subsequently blocked with 5% PBS/BSA for 1 hour at Room Temperature (RT) and incubated with the following primary antibodies used in pairs to detect the interactions: (I) rabbit anti-hTERT/mouse anti-TFAM, (ab119684; Abcam, Waltham, MA, USA); (II) rabbit anti-hTERT/mouse mtRPOL, (III) rabbit anti-hTERT/mouse anti-TWINKLE; (IV) rabbit anti-hTERT/mouse anti-TFB2M, in 5% PBS/BSA. After being washed with PBS, cells were incubated with secondary anti-rabbit antibody conjugated to Alexa 488 and secondary anti-mouse antibody conjugated to Alexa 568 (Thermo Fisher Scientific, Waltham, MA, USA). After washing in PBS, coverslips were mounted in ProLong Gold Antifade reagent (Invitrogen, Thermo Fisher Scientific, USA). All the samples were used to perform the Förster’s resonance energy transfer (FRET) by Fluorescence-lifetime imaging microscopy (FLIM). TERT was used as a FRET donor in all experiments and the Alexa 488 lifetime was measured with inverted SP8 Leica confocal microscope (Manheim, Germany) equipped with a single-molecule detection (SMD) module based on a Picoquant hardware solution (Berlin,Germany), by using a 470nm pulsed laser with a 40 MHz repetition rate and 63X oil immersion objective (N.A. 1.4). Lifetime values in each individual cell were determined by fitting the fluorescence decay with the built-in Symphotime Operation software (Picoquant, Germany). This was performed by integrating the signal from the pixels in a region of interest (ROI) with a single exponential model. To calculate ΔLifetime values, the mean lifetime of the cells in the donor-only condition (TERT-488) was calculated and then used to normalize data in all the analysed conditions and for each independent experiment.

To study the colocalization between LC3B and mitochondria, U2OS and U2OS-TERT-HA cell lines were transfected with the pCMV plasmids described above. 48 hours after transfection, cells were treated with MitoTracker DeepRed (Thermo Fisher Scientific) at the final concentration of 20nM for 30 minutes at 37°C. Immediately after staining, cells were processed by FRET/FLIM. Aquamarine was used as a FRET donor in all experiments and excited at 440nm pulsed laser. The microscopy and the calculation of ΔLifetime values were described above. Fluorescence co-localization between Aqua-LC3B puncta and MitoTracker Deep Red was calculated with the JaCoP plugin (Bolte and Cordelières, 2006).

To assess mitochondrial morphology, 5×10^4^ of U2OS and U2OS-TERT-HA cell lines were seeded on 15mm round coverslips placed onto 24-well plates. The immunofluorescence was performed as described above and the following primary antibody were used: mouse anti-TFAM and rabbit anti-PMPCB (16064-1-AP; Proteintech, Rosemont, IL, USA) in 5% PBS/BSA. After being washed with PBS, cells were incubated with secondary anti-mouse antibody conjugated to Alexa 568 and anti-rabbit antibody conjugated to Alexa 488. After washing in PBS, coverslips were mounted in ProLong Gold Antifade reagent. Images were acquired by an inverted SP8 Leica confocal microscope (Leica Microsystem, Wetzlar, Germany) using a 488nm laser at 0.25 % power and a 561 nm laser at 2% power and a 63X oil immersion objective (N.A. 1.4). Mitochondrial length and branching were calculated from confocal images on a total of 10 cells per condition and per biological replicate using ImageJ software as in Koopman et al. (2005).

### 2.4. Protein lysate preparation and Western Blot

Cells were lysed in RIPA buffer (150 mM NaCl, 1% Triton X-100, 0.5% DOC, 0.1% SDS, 50mM Tris-HCl pH 8.0), complemented with a protease inhibitor cocktail (Roche, Basel, Switzerland). Protein lysates (20μg) were loaded on SDS-PAGE and transferred onto a polyvinylidene fluoride (PVDF) membrane (Immobilion-P, Millipore, Merck, Darmstadt, Germany). After blocking in 3% bovine serum albumin (BSA) and 0.1% Tween, diluted in Tris-buffered saline solution (TBS), membranes were incubated with the following primary antibodies: mouse anti-β-actin, mouse anti-TFAM, rabbit anti-LC3B (Merck, Germany), mouse anti-Drp1 (611112 BD Pharmingen, San Diego, CA, USA) and mouse MitoBiogenesis Western Blot cocktail (Abcam, USA).

Finally, membranes were incubated with the appropriate HRP-conjugated secondary antibody (Bio-Rad Laboratories, Hercules, CA, USA). Proteins were visualized using Clarity Western ECL substrates (Bio-Rad, USA). Images were acquired on a ChemiDoc Imaging system (Bio-Rad, USA) and protein levels were quantified using the Image Lab software (Bio-Rad, USA). Experiments were repeated at least three times.

### 2.5. Quantification of mRNA levels by RT-qPCR

Total RNA was extracted with EuroGold TriFast reagent (Euroclone) and precipitated with 2-Propanol. RNA was reverse transcribed with the SuperScript™ III Reverse Transcriptase kit (Thermo Fisher Scientific), using random hexamers as primers for the reaction. Quantitative PCR was performed with SsoAdvanced Universal SYBR Green Supermix (Bio-Rad, Hercules, CA, USA) using the Agilent AriaMax real-time PCR system (Agilent Technologies, Santa Clara, CA, USA), with the following primers: TFAM-fw (AGCTCAGAACCCAGATGC); TFAM-rv (CCACTCCGCCCTATAAGC); GAPDH-fw (AGCCACATCGCTCAGACAC); GAPDH-rv (GCCCAATACGACCAAATCC).

Relative gene expression was calculated using the 2^-ΔΔCt^ method. The expression level of the target gene was compared to GAPDH in each cell line (using the difference in threshold cycles, or ΔCt), and then these values were normalized to the U2OS cell line. Each experiment was conducted at least in triplicate.

## 3. Results

### 3.1. TERT interacts with mitochondrial nucleoid complex proteins

We have previously shown that TERT is able to associate with mtDNA, affecting both replication and transcription (Marinaccio et al., 2025). However, previous results showed that this could be an indirect association, potentially due to the binding of TERT with mitochondrial nucleoid complex proteins (Marinaccio et al., 2025). To verify this hypothesis, we performed *in situ* Proximity Ligation Assay (*is*PLA) and Förster’s Resonance Energy Transfer (FRET) by Fluorescence Lifetime Imaging Microscopy (FLIM) between TERT and the mitochondrial transcription factor A (TFAM), mitochondrial polymerase (POLRMT), the helicase TWINKLE and the mitochondrial transcription factor B2 (TFB2M). These two techniques allow the identification of interactions between molecules that are at nanometre-scale distances. In particular, using the *is*PLA technique, interactions can be detected between proteins separated by distances of up to 40 nm, while FRET enables the detection of interactions of less than 10 nm (Hegazy et al., 2020; Lakowicz, 2006). We first used *is*PLA in U2OS-TERT-HA (U2OS cell line transduced with TERT-HA gene) and HCT-116 cell lines (Figure 1A and S1A-S1B) to analyse the binding between TERT and the above-mentioned mitochondrial nucleoid complex proteins. In the U2OS-TERT-HA cell line (Figure 1B), we observed a statistically significant difference between the number of *is*PLA-positive puncta in negative controls - samples in which only the anti-hTERT primary antibody was incubated - and those where TERT-TFAM were co-stained. These results were corroborated in HCT-116 cell line, which expresses TERT at physiological levels. In this model as well, we observed a significant difference between negative controls and the TERT-TFAM pair (Figure S1C). Subsequently, we verified these results by performing FRET/FLIM analyses (Figure 1C) in U2OS-TERT-HA cells. FRET/FLIM confirmed the interaction between TERT and TFAM (Figure 1D). In conclusion, we demonstrated that TERT is in spatial proximity with mitochondrial nucleoid proteins, and predominantly with TFAM.

**Fig. 1.**
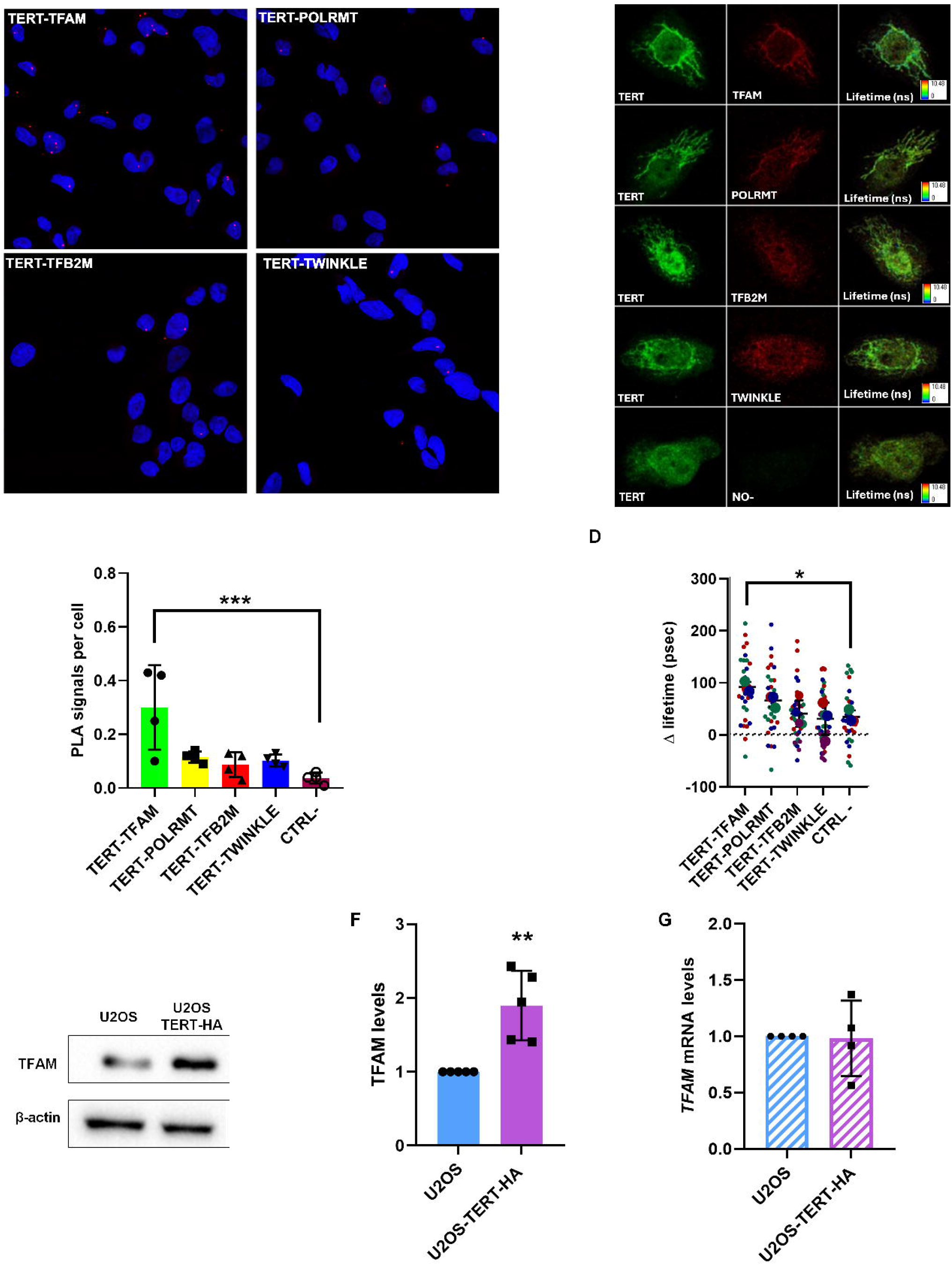
TERT interacts with TFAM. (A) Representative Proximity Ligation Assay images between TERT and four proteins of mitochondrial nucleoid complex. (B) *is*PLA signals per cell in U2OS-TERT-HA. Statistical analysis is performed comparing each sample to the CTRL- (sample in which only the anti-TERT primary antibody was incubated). Each dot corresponds to an independent biological replicate. (C) Representative fluorescence and lifetime images from FRET/FLIM analysis of U2OS-TERT-HA cells showing TERT (the FRET donor), four mitochondrial nucleoid complex proteins (FRET acceptors) and one donor-only condition which represents the internal control that lacks the acceptor and therefore cannot perform FRET. (D) Corresponding Δlifetime quantifications. A higher Δlifetime reflects a closer proximity between the donor and acceptor moieties. Statistical analysis is performed by comparing each sample to the CTRL- (donor-only condition); n = 10 cells per condition (small dots) in each of the experimental replicates. Large dots indicate mean values per replicate. (E) Representative Western blot of TFAM in U2OS and U2OS-TERT-HA cells. β-actin was used as a normalization control. (F) Quantification of TFAM protein levels in U2OS-TERT-HA cells, normalized to U2OS cells. Each dot corresponds to an independent biological replicate. (G) RT-qPCR evaluation of *TFAM* mRNA expression levels in U2OS-TERT-HA normalized to that in U2OS cells. Data are expressed as means ± SD \**P*<0.05; ** *P*<0.01; *** *P*<0.001 by one-way ANOVA (B-D) and unpaired t test (F-G).

To investigate whether TERT overexpression in U2OS alters TFAM expression, Western Blotting analysis of TFAM levels was carried out (Figure 1E), showing a significant increase in TFAM protein levels in the U2OS-TERT-HA cell line compared to U2OS (Figure 1F). On the other hand, *TFAM* mRNA levels were not significantly different between U2OS-TERT-HA and U2OS cells (Figure 1G). These findings show that TERT overexpression increases TFAM protein abundance, while leaving its transcript levels unaltered.

### 3.2. TERT induces mitochondrial fission in a TFAM-independent manner

We previously observed that TERT-overexpressing cells show a higher number of mitochondria per cell and with a smaller area than those in controls (Marinaccio et al., 2025). We therefore asked whether TERT overexpression induces mitochondrial fission and whether the potential induction may be TFAM-dependent. Mitochondria were stained for the matrix protein PMPCB, and mitochondrial morphology analyses were performed by analysing mitochondrial length, branching and the number of mitochondria (Koopman et al., 2005) in controls and U2OS-TERT-HA cells. These analyses were then performed in the presence or absence of *TFAM* silencing to evaluate the functional relevance of the TERT/TFAM interaction towards mitochondrial morphology (Figure 2A and S2A-B). We observed a significant reduction in mitochondrial length in U2OS-TERT-HA SCR (negative control siRNA) compared to U2OS SCR cells (Figure 2B-I), confirming previous results obtained with Transmission Electron Microscopy (TEM) (Marinaccio et al., 2025). However, the difference between U2OS and U2OS-TERT-HA cells was abolished upon *TFAM* downregulation. Of note, no differences in mitochondrial branching and in the number of mitochondria per cell were found in all samples analysed (Figure 2BII and S2C).

**Fig. 2.**
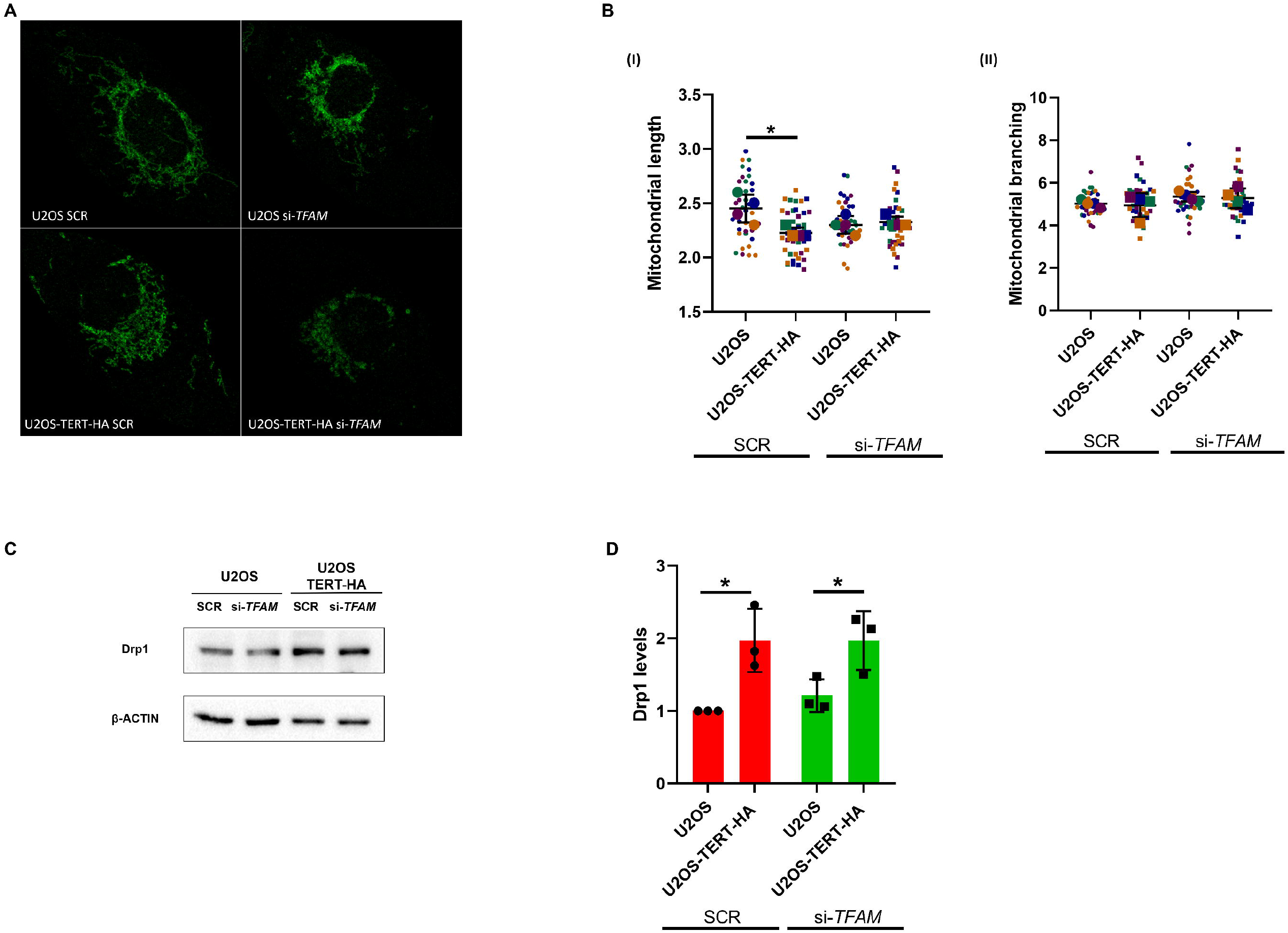
TERT induces mitochondrial fission regardless of *TFAM* downregulation. (A) Representative fluorescence images of U2OS and TERT-HA overexpressing cell line showing mitochondria labelled with an anti-PMPCB antibody, in the presence or absence of si-*TFAM*. (B) The graphs show the analysis of mitochondrial length (I) and branching (II), with and without si-*TFAM. n* = 10 cells per condition (small dots) in each of three experimental replicates. Large dots indicate mean values per each replicate. (C) Representative Western blot of Drp1 in U2OS and U2OS-TERT-HA cell lines with and without si-*TFAM*. β-actin was used as control protein. (D) Quantification of Drp1 protein levels in U2OS and U2OS-TERT-HA cell lines with and without si-*TFAM*, normalized to the U2OS SCR samples. Each dot corresponds to an independent biological replicate. Data are expressed as means values ± SD. Statistical analysis is performed by comparing each sample to U2OS cell line \**P*<0.05 by two-way ANOVA with Tukey’s (B) and Sidak’s (D) multiple comparisons tests. Scale bar: 10 µm.

To determine whether the observed mitochondrial fragmentation was caused by accelerated mitochondrial fission, we analysed the protein levels of Drp1, the main regulator of this process (Di Nottia et al., 2021). Drp1 abundance was analysed in U2OS and U2OS-TERT-HA cells with and without si-*TFAM* (Figure 2C). U2OS-TERT-HA cells showed higher levels of Drp1 compared to U2OS cells, and this increase occurs independently of the presence or absence of TFAM (Figure 2D). In conclusion, TERT enhances mitochondrial fragmentation and organelle fission regardless of its interaction with TFAM.

### 3.3. The TERT/TFAM interaction promotes auto/mitophagy activation

The increase in auto/mitophagosomes number previously observed with TEM in TERT-overexpressing cells (Marinaccio et al., 2025) and the induction of mitochondrial fission may be indicative of ongoing mitochondrial turnover by mitophagy. This process would generate small mitochondrial fragments targeted for degradation through the autophagy pathway. Therefore, we investigated the activation of auto/mitophagy using the LC3B biosensor, a FRET-based molecular probe responding to the priming of LC3B by ATG4B (autophagy related 4B cysteine peptidase) (Gökerküçük et al., 2024). FRET/FLIM analyses were performed in normal and TERT-overexpressing U2OS cells, in the presence or absence of TFAM (Figure 3A). U2OS-TERT-HA cells showed a statistically significant decrease of the LC3B biosensor Δlifetime compared to that observed in U2OS cells, indicating a more pronounced activation of auto/mitophagy in cells expressing TERT compared to controls (Figure 3B). A version of the LC3B biosensor uncleavable by the ATG4B protease was used as a control for the maximal FRET effect detectable with this probe and showed no difference in FRET/FLIM values in the two cell lines (Figure S3A-B).

**Fig. 3.**
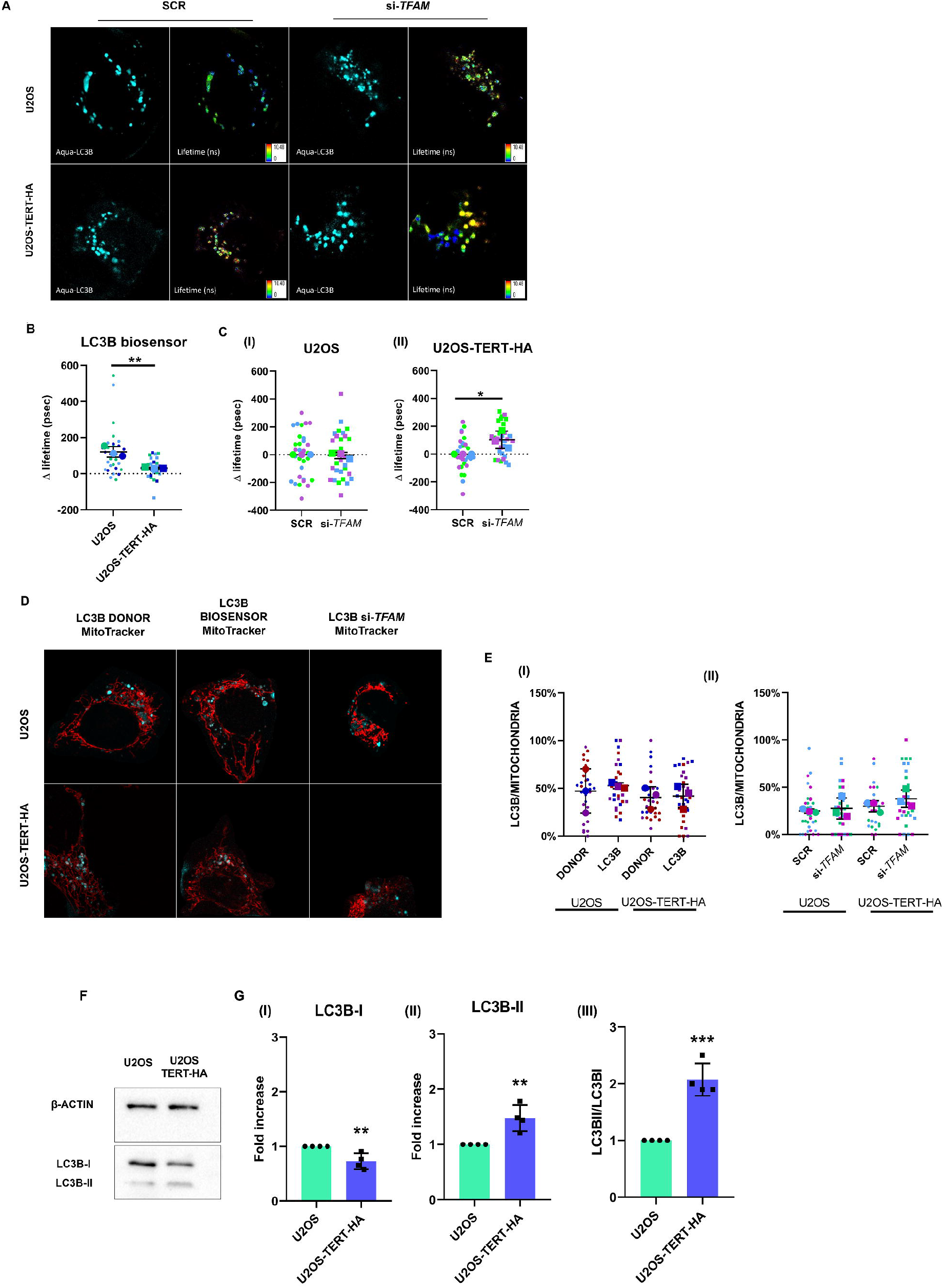
TERT promotes auto/mitophagy activation thanks to the binding with TFAM. (A) Representative fluorescence and lifetime images of U2OS and U2OS-TERT-HA cells showing the LC3B biosensor with and without si-*TFAM* respectively. (B) Quantification of the number of Aqua-LC3B-II puncta in cells expressing the LC3B biosensor in U2OS and U2OS-TERT-HA cell lines. (C) Quantification of the number of Aqua-LC3B-II puncta upon *TFAM* downregulation compared to controls. ΔLifetime quantifications (in psec) performed in U2OS (I) and TERT-HA transfected cells (II) with and without si-*TFAM*. n = 10 cells per condition (small dots) in each of three experimental replicates. Large dots indicate mean values per replicate. (D) Representative fluorescence images showing the object-based colocalization between MitoTracker (pseudocolored red) and LC3B-II, (pseudocolored cyan) in U2OS and U2OS-TERT-HA cells transfected with the Aqua-LC3B-II biosensor with and without si-*TFAM*. (E) Quantification of Aqua-LC3B-II puncta structures colocalizing with MitoTracker Red in cells expressing the LC3B biosensor with (II) and without (I) si-*TFAM. n* = 10 cells per condition (small dots) in each of three experimental replicates. Large dots indicate mean values for each replicate. (F) Representative western blot images of LC3B-I and LC3B-II in U2OS and U2OS-TERT-HA cells. β-actin was used as a control protein. (G) Quantification of LC3B-I (I), LC3B-II (II) and LC3B-II/LC3B-I (III) protein levels in the U2OS-TERT-HA cell line, normalized to those in the U2OS cell line. Each dot corresponds to an independent biological replicate. Data are expressed as means ± SD \**P*<0.05 ** *P*<0.01 *** *P*<0.001 by unpaired t-test (B-C-G) and comparisons were not significant after by two-way ANOVA (E).

We then asked whether TFAM was necessary for the role of TERT in activating auto/mitophagy. To address this question, we used the LC3B biosensor upon *TFAM* downregulation in the two cell types (Figure 3A). In U2OS cells, auto/mitophagy activation remained unchanged despite *TFAM* downregulation (Figure 3CI). In contrast, *TFAM* knockdown induced a significant increase in the Δlifetime of the LC3B biosensor in U2OS-TERT-HA cells compared to SCR controls (Figure 3CII). This result indicates a decrease of auto/mitophagy activation when *TFAM* is depleted. This was not due to a differential localization of the sensor at mitochondria, as the colocalization between the LC3B biosensor and the MitoTracker DeepRed dye was constant across conditions and cell lines (Figure 3D-E). Furthermore, colocalization analysis revealed an overlap of approximately 50% of LC3B-positive objects with mitochondria. These findings suggest that the autophagic response observed in our model may involve both mitophagy and bulk autophagy (Figure 3EI).

We then analysed the activation of LC3B by Western blotting (Figure 3F). We observed a reduction in the inactive form of LC3B – LC3B-I – (Figure 3GI) in U2OS-TERT-HA cells, associated with an increase in the active one – LC3B-II – (Figure 3GII), thereby leading to an elevated LC3BII/LC3BI ratio (Figure 3GIII). This orthogonal approach confirmed that TERT overexpression promotes auto/mitophagy activation.

In conclusion, these findings show that TERT activates auto/mitophagy through its interaction with TFAM without changing the proximity between LC3-positive autophagosomes and mitochondria.

### 3.4. TERT expression induces a remodelling of the respiratory chain in a TFAM-dependent manner

Mitobiogenesis and mitophagy are two interdependent processes that must be under tight control (Cardoso-Pires and Vieira, 2024). Taking into account the role of TERT in mitochondrial fission and the TERT/TFAM-related activation of auto/mitophagy, we evaluated whether this functional interaction may also regulate mitochondrial biogenesis. To this end, we used the Mitobiogenesis Western Blotting cocktail, which consists of antibodies targeting two subunits of different oxidative phosphorylation enzyme complexes, such as the nuclear-encoded Complex II protein SDH-A and the mitochondrially-encoded Complex IV subunit COX-I. Western blotting analyses were performed in U2OS and U2OS-TERT-HA cells, with and without *TFAM* downregulation (Figure 4A). We observed a significant decrease in COX-I protein levels in U2OS-TERT-HA compared to U2OS SCR. Instead, the SDH-A levels remained unaltered in the two cell lines (Figure 4A-B). We also observed that the TERT-dependent lowered abundance of COX-I was abolished in the absence of *TFAM*. Indeed, no significant difference in COX-I protein levels was detected in U2OS cells compared to U2OS-TERT-HA when *TFAM* is silenced (Figure 4A-B). Therefore, a statistically significant decrease in the COX-I/SDHA ratio was observed in U2OS-TERT-HA SCR compared with U2OS SCR, whereas this reduction was not observed in the TFAM-depleted samples (Figure 4C).

**Fig. 4.**
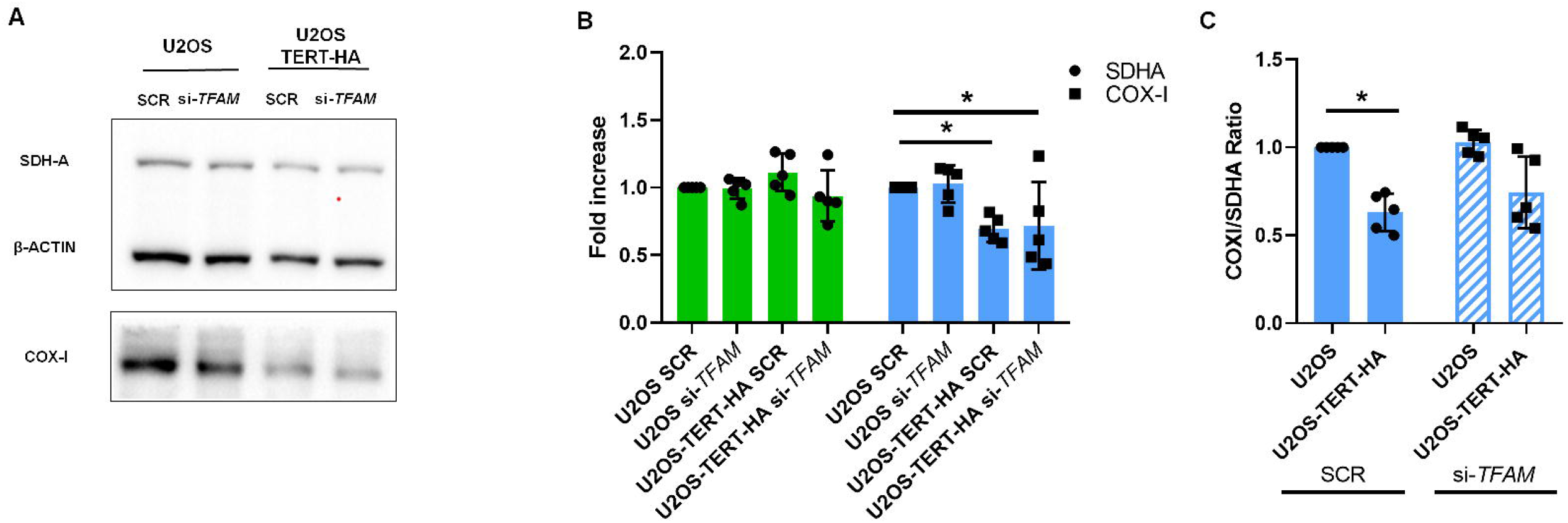
TERT induces a remodelling of the respiratory chain. (A) Representative Western blotting images of SDH-A and COX-I in U2OS and U2OS-TERT-HA cell lines with and without si-*TFAM*. β-actin was used as a loading control. (B) Quantification of the SDH-A and COX-I protein levels in U2OS and U2OS-TERT-HA cell lines with and without si-*TFAM*, normalized to U2OS SCR. (C) Quantification of the COX-I/SDH-A ratio in U2OS and TERT-HA overexpressing cells with and without *TFAM* downregulation, normalized to U2OS SCR. Each dot corresponds to an independent biological replicate. Data are expressed as means ± SD. \**P*<0.05 by Two-way ANOVA with Sidak’s multiple comparisons test.

In conclusion, our results suggest that TERT plays a role in the remodelling of the mitochondrially-encoded subunit COX-I rather than playing a broader role in mitochondrial biogenesis. This is further supported by the previously reported association between TERT and the COXI mtDNA region (Marinaccio et al., 2025).

## 4. Discussion

In our previous work (Marinaccio et al., 2025), we showed that TERT is able to associate with different mitochondrial genes, in agreement with Sharma et al. (2012). Moreover, we observed that TERT affects both mitochondrial replication and transcription (Marinaccio et al., 2025). However, whether the association of TERT with mtDNA occurred through direct binding or an indirect interaction remained unknown. In this work, we demonstrated that TERT predominantly binds TFAM using a combination of quantitative fluorescence microscopy and biochemical approaches in U2OS-TERT-HA cells, thereby supporting an indirect binding of TERT with mtDNA. The same association was also observed in HCT-116 cells. In both cell lines though, the extent of *is*PLA signals per cell remains relatively low, suggesting that TERT and TFAM do not form a stable complex. However, the TERT/TFAM interaction is the strongest among those involving TERT and POLRMT, TFB2M or TWINKLE. Since they are components of the mtDNA replication and transcription machinery (Falkenberg, 2018), their interaction is likely to be transient, occurring only during specific stages.

*is*PLA allows the detection of interactions between proteins separated by distances of up to 40 nm. To further validate these findings, we subsequently performed FRET/FLIM analyses, which provide a more direct assessment of molecular interactions occurring within nanometer-scale distances (less than 10 nm) (Hegazy et al., 2020; Lakowicz, 2006). The application of a more sensitive and stringent experimental approach further confirmed that TFAM is the main protein interacting with TERT among those screened with *is*PLA. Collectively, these findings provide the first evidence that TERT physically interacts with TFAM within the mitochondrial matrix. Rather than directly binding mtDNA, TERT appears to associate with mtDNA indirectly through interaction with TFAM, a protein involved in mitochondrial DNA replication and transcription. This observation would explain the results obtained in our previous work, in which we demonstrated that TERT promotes both mtDNA replication and mitochondrial transcription (Marinaccio et al., 2025).

Using an in vitro pull-down assay, Polanská et al. (2012) reported a direct interaction between human TERT and HMGB1. HMGB1 is a chromatin-associated protein functioning as a DNA chaperon in transcription, replication, recombination and repair (Agresti and Bianchi, 2003). Specifically, HMGB1 binds to the RID2 domain of TERT. Since TFAM belongs to the high-mobility group box (HMGB) protein family as HMGB1 and contains two HMG-box domains (Yang et al., 2025), it is tempting to speculate that TERT and TFAM interaction can occur within the RID2 and HMG domains. Future evidence will determine the protein domain(s) responsible for the TERT/TFAM interaction. TFAM is bound to mtDNA and it is essential for its synthesis, expression and packaging (Asin-Cayuela and Gustafsson, 2007). Lu et al. (2013) showed that TFAM is phosphorylated by cAMP-dependent protein kinase (PKA) within its HMGB1 domain. This phosphorylation reduces the ability of TFAM to bind mitochondrial DNA, with the consequence of creating a pool of TFAM free from DNA. This pool is recognized and degraded by the mitochondrial Lon protease. Reducing Lon levels in cells with severe mtDNA depletion rates increases both TFAM protein levels and mtDNA content. In our work, we observed that TERT-overexpressing cells show higher levels of the TFAM protein than controls, and this without changes in the mRNA abundance. This result suggests that TERT may act to stabilize TFAM at the protein level, potentially by decreasing its degradation rather than regulating its transcriptional levels. TERT may elicit a similar effect as Lon deficiency although this is not accompanied by an increase in mtDNA copy number in TERT-HA cells (Marinaccio et al.,2025). This may indicate that the balance between stabilization and degradation is subtle in this model, with small changes allowing TFAM stabilization.

Mitochondria are an interconnected network tightly governed by a dynamic equilibrium between fission and fusion. Mitochondrial fission promotes the segregation of damaged mitochondria which are subsequently eliminated through mitophagy (Di Nottia et al., 2021). The main protein regulating mitochondrial fission is Drp1, a GTP-hydrolyzing enzyme which oligomerises into filaments around the outer mitochondrial membrane, forming a ring-like structure that constricts the membrane until fission occurs (Mears et al., 2011). Using transmission electron microscopy (TEM), we previously showed that there is an increase in the number of mitochondria upon TERT expression, and that mitochondria are smaller than in control conditions. This is accompanied by an increased amount of auto/mitophagosomes. Mitochondrial fragmentation together with the presence of auto/mitophagosomes hinted at the activation of mitochondrial turnover (Marinaccio et al., 2025). Building on these results, an important question that remained to be addressed was whether TERT directly triggers these processes. Using confocal microscopy, we here confirmed that there is a reduced mitochondrial length in TERT-overexpressing cells compared to control cells. To determine whether this lowered mitochondrial length was due to an increased mitochondrial fission, we analysed the levels of Drp1. Drp1 protein abundance was higher in TERT-overexpressing cells compared to U2OS, and this occurred in a TFAM-independent manner. The independence of this process from TFAM may be due to the different localization of TFAM and Drp1 in mitochondria. With TFAM in the matrix and Drp1 on the outer mitochondrial membrane (OMM) it is conceivable that the absence of the matrix partner of TERT does not impact the ability of Drp1 to be recruited at the OMM and perform its function in TERT-overexpressing cells.

The increase in mitochondrial fission demonstrated above may enhance mitophagy by generating smaller mitochondrial fragments that can be targeted for degradation. To test this hypothesis and considering that TERT overexpression induced an increase in auto/mitophagosomes (Marinaccio et al., 2025), we used the LC3B FRET biosensor to visualize ongoing auto/mitophagy events. LC3B is normally present in an inactive state as LC3B-I, carrying five amino acid residues at its C-terminus. For LC3B to become functionally active, these residues must be cleaved by ATG4B, obtaining LC3B-II. This processing step is required for the conjugation of LC3B-II to the phagophore and thus initiates autophagy (Agrotis et al., 2019). The LC3B biosensor is tagged with two different fluorophores at the N- and C-termini of the uncleaved, full-lenght LC3B protein. When ATG4B is inactive and unable to prime LC3B-I, the fluorophores remain in close proximity and FRET occurs. Conversely, when ATG4B is active and cleaves the C-terminal region of LC3B, the protein becomes active and capable of initiating the autophagy process, resulting in a loss of FRET (Gökerküçük et al., 2024). LC3 biosensor analysis was performed in combination with MitoTracker staining to determine whether the autophagic component detected by the biosensor could be attributed, at least in part, to mitophagy. To further investigate this possibility, colocalization analysis revealed approximately 50% overlap of LC3B-positive objects with mitochondria, thereby confirming the contribution of mitophagy to the observed autophagic process. Using this approach, we found that TERT-overexpressing cells display enhanced ongoing auto/mitophagy compared to control cells where this process is partially stalled or slowed down. We showed that this phenomenon depends on the presence of TFAM, since its downregulation phenocopies auto/mitophagy in control cells. Overall, these data indicate that TERT and TFAM cooperate to enhance auto/mitophagy flux activation. TERT and TFAM are located in the matrix and in proximity to the IMM (Farge and Falkenberg, 2019; Haendeler et al., 2009), hence leading to the intriguing hypothesis that the autophagy receptor of LC3B may be located at this compartment as well. So far, Prohibitin-2 (PHB2) is the only IMM receptor of LC3B during mitophagy (Wei et al., 2017). Future results will determine whether PHB2 is also an interactor of the TERT/TFAM pair and required to complete mitochondrial clearance in this paradigm. Another compelling result that supports these findings was obtained by analysing the two forms of LC3B. In TERT-overexpressing cells, we observed a decrease in the inactive LC3B-I form accompanied by a corresponding increase in the active LC3B-II form, resulting in an elevated LC3B-II/LC3B-I ratio. Since an increased LC3B-II/LC3B-I ratio is a well-established hallmark of autophagosome formation and enhanced autophagic activity during mitophagy (McLeland et al., 2011), these results further support the increased activation of the auto/mitophagy process when TERT is present.

An involvement of TERT in mitophagy was observed by Shin and Chung (2020), who demonstrated that human TERT increases mitophagy by modulating PINK1 abundance. The PINK1/Parkin pathway is one of the major mitophagy pathways by promoting the recognition and clearance of damaged mitochondria (Narendra et al., 2010). Under physiological conditions, PINK1 is imported into mitochondria and cleaved by mitochondrial processing peptidase (MPP) in the matrix and by PARL in the inner mitochondrial membrane (Greene et al., 2012; Jin et al., 2010). TERT binds directly to the catalytic β subunit of MPP, inhibiting the proteolytic processing of PINK1 and resulting in the accumulation of its full-length. At this point, PINK1 becomes activated, resulting in the recruitment of Parkin and autophagy receptor proteins localized on the surface of the OMM, which initiate the formation of the autophagosome. This process leads to the disruption of the outer mitochondrial membrane, exposing PHB2, an inner mitochondrial membrane protein that can interact with LC3B and thereby enhance the efficiency of Parkin-dependent mitochondrial degradation (Uoselis et al., 2023). The recent report of Yang et al. (2026)instead showed that the overexpression of TOMM20 enhances mitophagy by promoting the mitochondrial translocation of TERT in an in vitro model of membranous nephropathy. Such enhanced mitophagy is associated with increased PINK1 expression, a higher LC3-II/LC3-I ratio, decreased p62 levels, and increased mitochondrial Parkin recruitment. The study suggests that TOMM20-mediated regulation of mitophagy requires TERT, as TERT downregulation abolished mitophagy. Although in a different paradigm, this study supports our own results, showing that TERT depletion restores normal auto/mitophagy rates. Still, more extensive analyses will establish whether TERT- and TFAM-dependent mitophagy always depends on PINK1 and Parkin, or whether such dependency is limited to these specific pathophysiological conditions. Altogether, the evidence reported in these studies further supports our findings which showed that TERT promotes mitochondrial fragmentation and mitochondrial fragmentation by mitophagy. While these studies focus on TERT acting as a positive regulator of mitophagy under stress or damaged conditions, we here show that TERT overexpression is sufficient to trigger these mitochondrial events.

Biogenesis and mitophagy are two interdependent processes (Cardoso-Pires and Vieira, 2024), but the increase in auto/mitophagy activation is not correlated to an increase in biogenesis when TERT is overexpressed. Under these conditions, the observed reduction in the mitochondrially-encoded COX-I protein levels may reflect an alteration in cellular metabolism, potentially compensated by the increased abundance of the nuclear-encoded SDHA protein. Recently, Kugapreethan et al. (2025), demonstrated that SDHA accumulates during dysfunctions of Complexes II, III and IV. Thereby, the assembly of respiratory chain complex can adapt to different energetic requirements of the cell under stress. This observation agrees with our previous findings (Marinaccio et al., 2025), which demonstrated reduced mitochondrial ATP production and a metabolic shift toward glycolysis in TERT-overexpressing cells.

In conclusion, we provide the first evidence that TERT physically interacts with TFAM within the mitochondrial matrix, supporting an indirect association of TERT with mtDNA and identifying the TERT/TFAM association as a potential molecular mechanism underlying TERT-mediated regulation of mitochondrial homeostasis. The interaction is associated with increased mitochondrial fission and enhanced auto/mitophagy. This could suggest that TERT promotes the turnover of dysfunctional mitochondria and may thereby contribute to the maintenance of mitochondrial and cellular fitness. Together with our previous demonstration (Marinaccio et al., 2025) that TERT affects mtDNA replication and transcription, these findings extend the role of TERT beyond its canonical function in telomere maintenance and support its involvement in the preservation and renewal of the mitochondrial network. The identification of the TERT/TFAM interaction provides a mechanistic framework for understanding how TERT may influence mitochondrial quality control and cellular adaptation to stress. This may be particularly relevant to mitochondrial and neurodegenerative disorders, in which mitochondrial dysfunction and oxidative stress are central features. Conversely, the same pathway may have different implications in pathological contexts such as cancer, in which limiting the maintenance and clearance of damaged mitochondria could contribute to mitochondrial stress and cell death. Thus, defining the molecular mechanisms governing TERT-dependent mitochondrial quality control may provide important insights into how mitochondrial fitness is maintained in health and disease.

## CRediT authorship contribution statement

**Erica Rossi:** Data curation; Formal analysis; Investigation; Methodology; Validation; Visualization; Roles/Writing - original draft; Writing - review & editing.

**Jessica Marinaccio:** Data curation; Formal analysis; Investigation; Validation; Roles/Writing - original draft; Writing - review & editing

**Ilaria Festo:** Formal analysis; Investigation; Methodology; Validation.

**Ion Udroiu:** Conceptualization; Formal analysis; Writing - review & editing.

**Emanuela Micheli:** Conceptualization; Writing - review & editing.

**Giulia Bertolin:** Conceptualization; Funding acquisition; Methodology; Project administration; Resources; Writing - review & editing.

**Antonella Sgura:** Conceptualization; Funding acquisition; Project administration; Resources; Writing - review & editing.

## Acknowledgments

We thank Xavier Pinson and Stéphanie Dutertre at the Microscopy Rennes Imaging Center (MRic, Biologie, Santé, Innovation Technologique-BIOSIT, Rennes, France) for assistance with FLIM experiments. MRic is member of the national infrastructure France-BioImaging, supported by the French National Research Agency (ANR-10-INBS-04). We thank Chloé Bertin (IGDR) for technical assistance.

## Data availability

Source microscopy data are available on Zenodo (10.5281/zenodo.22825153). All other data are available from the corresponding authors (G.B., A.S.) upon request.

