## Supplementary material for "TERT interacts with TFAM to activate mitochondrial fragmentation and auto/mitophagy": Supplementary materials.pdf

### Figure S1

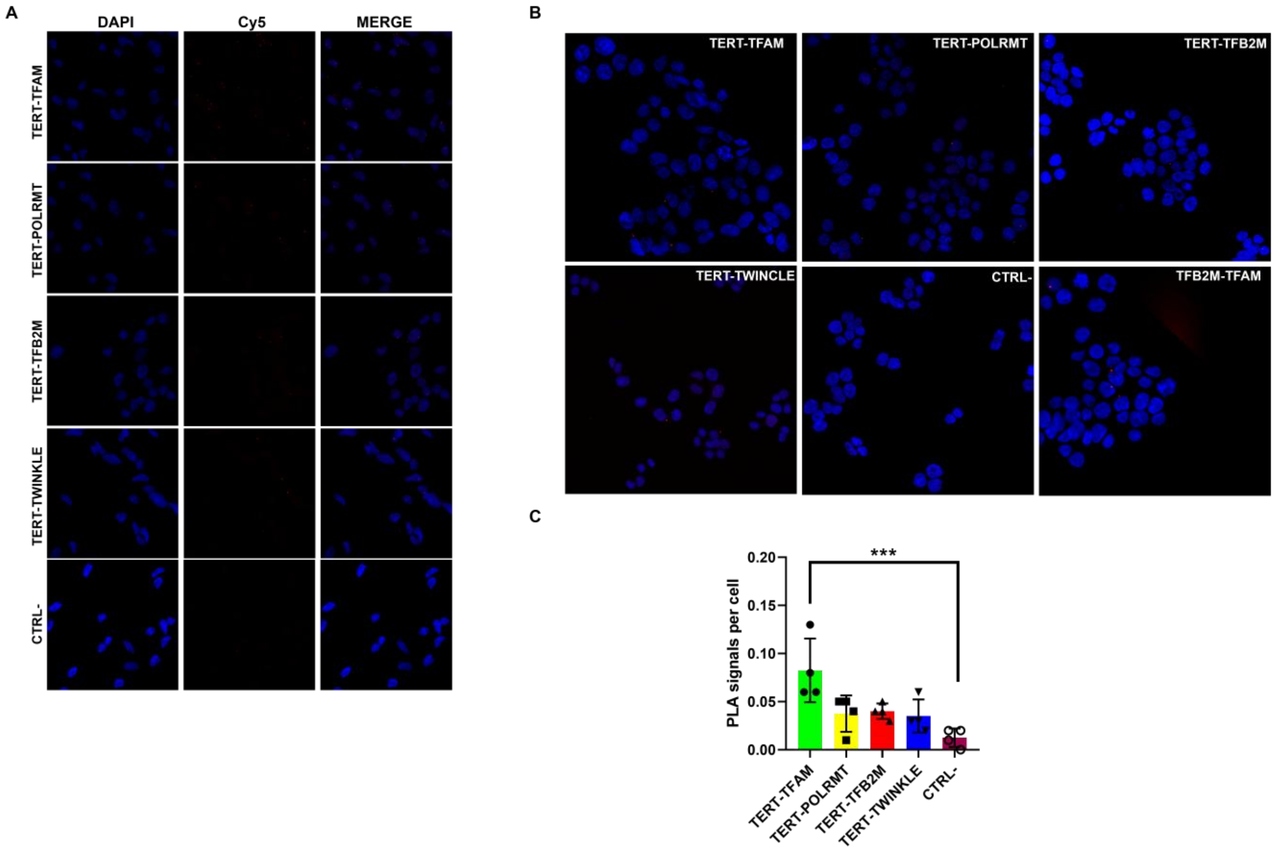


**Fig. S1. TERT interacts with TFAM in HCT-116 cell line.** (A) Representative *is*PLA images of U2OS-TERT-HA cells showing the proximity between TERT and TFAM, POLRMT, TFB2M, TWINKLE and a negative control sample (CTRL-), which refers to U2OS-TERT-HA incubated with the TERT antibody only. (B) Representative *is*PLA images of HCT-116 cells showing the interaction between TERT and TFAM, POLRMT, TFB2M, TWINKLE and negative control sample (CTRL-), which refers to HCT-116 incubated with the TERT antibody only. (C) *is*PLA signals were counted and normalized to cell number in each region in HCT-116 cell. Each dot corresponds to an independent biological replicate. Data are expressed as means ± SD. Statistical analyses were performed by comparing each sample to the CTRL- *** $P$<0.001 by one-way ANOVA.

### Figure S2

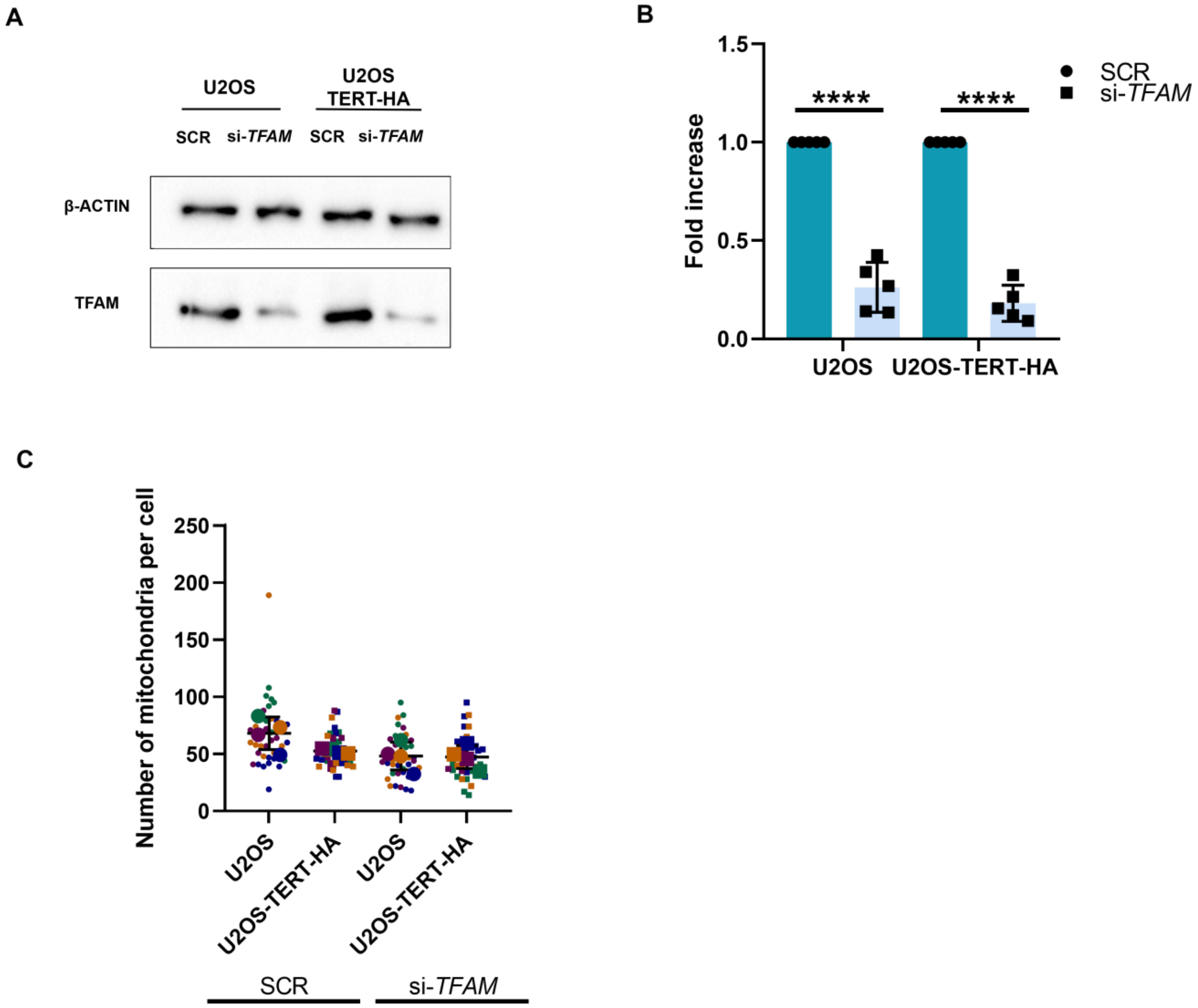


**Fig. S2. The number of mitochondria is independent of TERT and TFAM.** (A) Representative Western blot image of TFAM in U2OS and U2OS-TERT-HA cell lines with and without si-*TFAM*. β-actin was used as a loading control. (B) Quantification of TFAM protein levels in U2OS and U2OS-TERT-HA cell lines with si-*TFAM*, normalized to SCR samples of each cell line. Each dot corresponds to an independent biological replicate. (C) Analysis of the number of mitochondria with and without si-*TFAM*. n = 10 cells per condition (small dots) in each of three experimental replicates. Large dots indicate mean values for each replicate. Data are expressed as means ± SD. Statistical analyses were performed by comparing each sample to the SCR sample of each cell line. **** $P$<0.0001 by two-way ANOVA with Tukey's multiple comparisons test.

### Figure S3

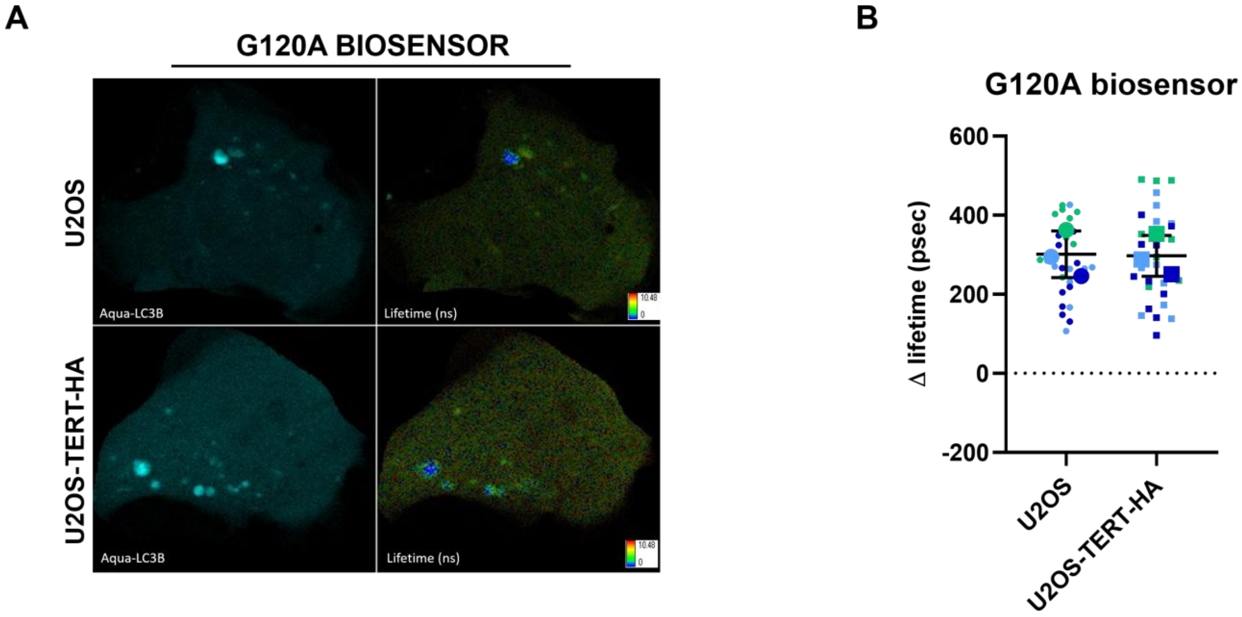


**Fig. S3. LC3B-G120A FRET positive control.** (A) Representative fluorescence and lifetime images of U2OS and U2OS-TERT-HA cells expressing the Aquamarine-LC3-G120A-TdLanYFP FRET positive control. (B) Quantification of the number of Aqua-LC3B-II puncta in cells expressing the G120A biosensor. ΔLifetime quantifications (in psec) performed in U2OS and U2OS-TERT-HA cells transfected as indicated. n = 10 cells per condition (small dots) in each of three experimental replicates. Large dots indicate mean values for each replicate. Data are expressed as means ± SD. Comparisons were not significant after by unpaired t-test.
